## Supplementary material for "A simplified method for CRISPR-Cas9 engineering of *Bacillus subtilis*": SI material

**Table S1: Strains used in the study.**

**Table S2: Oligos used in the study.**

**Fig S1: Detailed outline of CRISPR-Cas based editing strategy.**

**Fig S2: General strategy for assembly of repair template.**

**Fig S3: pAJS23 plasmid map and sequence (text) file**

### Table S1. Strains used in the study

| Strain | Genotype | Construction | Reference |
|---|---|---|---|
| *E. coli* | | | |
| ECE358 | *pJOE8999 (kan)* | | (1) |
| HEAS25630 | *pJOE8999-gRNA(erm)* | pAJS23→DH5α | This study |
| HEAS25655 | *pJOE8999-gRNA(erm)-prkC-Sau-rpe(repair)* | pAJS24→ DH5α | This study |
| HEAS25666 | *pJOE8999-gRNA(erm)-prkC-Sau-rpe(repair)* | pAJS24 → TG1 | This study |
| HEAS1215 | p*JOE8999-gRNA*(*erm*)-*yqgC-sodA* without S936 (Ref nt. position 2586043-2586220 deleted)-repair template | pAJS28→DH5α | This study |
| HEAS1223 | p*JOE8999-gRNA*(*erm*)-yqgC-sodA without S936 (Ref nt. position 2586043-2586220 deleted)-repair template | pAJS28→TG1 | This study |
| HEAS1262 | p*JOE8999-gRNA*(*erm*)-*yceF**-Ile206Thr(repair) | pAJS27→DH5α | This study |
| HEAS1258 | p*JOE8999-gRNA*(*erm*)-*yceF**-Ile206Thr(repair) | pAJS27→TG1 | This study |
| HEAS1183 | p*JOE8999-gRNA*(*erm*)-*yqgB-yqgC-gfp-sodA*(repair) | pAJS26→DH5α | This study |
| HEAS1185 | p*JOE8999-gRNA*(*erm*)-*yqgB-yqgC-gfp-sodA*(repair) | pAJS26→TG1 | This study |
| HEAS1214 | p*JOE8999-gRNA*(*erm*)-*yqgB-yqgC-sodA-FLAG-yqgE* (repair) | pAJS25→DH5α | This study |
| HEAS1222 | p*JOE8999-gRNA*(*erm*)-*yqgB-yqgC-sodA-FLAG-yqgE* (repair) | pAJS25→TG1 | This study |
| HEAS1206 | p*JOE8999-gRNA*(*erm*)-*yqgB*-scar-*sodA -yqgE* (repair)-0.5kb length | pAJS29→ DH5α | This study |
| HEAS1212 | p*JOE8999-gRNA*(*erm*)-*yqgB*-scar-*sodA -yqgE* (repair) )-0.5kb length | pAJS29→TG1 | This study |
| HEAS1206.1 | p*JOE8999-gRNA*(*erm*)-*yqgB*-scar-*sodA -yqgE* (repair)-1.5kb length | pAJS30→ DH5α | This study |
| HEAS1212.1 | p*JOE8999-gRNA*(*erm*)-*yqgB*-scar-*sodA -yqgE* (repair) )-1.5kb length | pAJS30→TG1 | This study |
| *B. subtilis* | | | |
| 168 | *trpC2* | Lab strain | Lab stock |
| CU1065 | *trpC2 att SPβ* (WT) | Lab strain | Lab stock |
| HB20401 | *trpC2 cpgA*::*erm* | BGSC | Lab stock |
| HBYL844 | *trpC2 cpgA*::*cpgA-Sau* | pAJS24$_{CRISPR}$→HB20401 | This study |
| HBYL264 | *trpC2 yqgC::erm* | *BGSC yqgC::erm* → CU1065 | This study |
| HBYL1239 | *trpC2* Δ2586043-2586220 intergenic deletion between *yqgC-sodA* | pAJS25$_{CRISPR}$→ HBYL264 | This study |
| HBYL344 | *trpC2 yceF*::*erm* | BGSC yceF::erm --> 168 | Lab stock |
| HBYL1260 | *trpC2 yceF* (Ile206Thr)* | pAJS26$_{CRISPR}$→ HBYL344 | This study |
| HBYL1246 | *trpC2 yqgC-gfp* | pAJS27$_{CRISPR}$→ HBYL264 | This study |
| HBYL1249 | *trpC2 sodA-FLAG* | pAJS28$_{CRISPR}$→ HBYL264 | This study |
| HBYL260 | *trpC2* Δ*yqgC* | PAJS29 $_{CRISPR}$→ HBYL264 | This study |
| HBYL261 | *trpC2* Δ*yqgC* | PAJS30 $_{CRISPR}$→ HBYL264 | This study |

### Table S2. Oligos used in the study

| Primer Name | Sequence | Reference |
|---|---|---|
| ermgRNAF | TACGTTTGAAATCGGCTCAGGAAA | This study |
| ermgRNAR | AAACTTTCCTGAGCCGATTTCAAA | This study |
| prkCFrepair | **AAGGCCAACGAGGCC**ACGGATCCTAAAGCGGATACCACAG | This study |
| LFHprkC-Sau-rsgA-R | GATTTCACTATTCGACCTGTCTTCAAATTTTCCCTCCTTGTTATTCATCTTTC | This study |
| LFHprkC-Sau-rsgA-F | GAAAGATGAATAACAAGGAGGGAAAATTTGAAGACAGGTCGAATAGTGAAATC | This study |
| Rpe-downrepair-R | **AAGGCCTTATTGGCC**TAGATCGGGAATGAGATTTTTCGGGCCTC | This study |
| yqgC1-seqF | CTTTTGTCATTCCGGTTGCAGGGATC | This study |
| gfp1-seqR | AAGTTTTCCGTATGTTGCATCACCTTCAC | This study |
| 0.5kb-yqgC-F | **AAGGCCAACGAGGCC**CCCTGTACTTACATAATAAG | This study |
| 0.5kb-yqgC-R | TTGGCCAATAAGGCCTCACGTATGTGTTGTGGTGTTTC | This study |
| 0.725kb-yqgC-up-F | **AAGGCCAACGAGGCC**AATAAGCAGAAAGCTTCCAGAGCTG | This study |
| GFP-LFHF | ATGATCGGTTACTTTTTATGGACGGTCCTACGTAAAGGAGAAGAACTTTTCACTGGAGTT | This study |
| GFP-LFHR | CATTAAACCTGCCGCCAGCATTTCTTATTTGTATAGTTCATCCATGCCATGTGTAATC | This study |
| 1.463kb-yqgC-R | **AAGGCCTTATTGGCC**CACGACAGCCAGGGCAAATAAACTG | This study |
| 1.028kb-yqgC-F | **AAGGCCAACGAGGCC**TATTGTCCAGAGCTTGTTC | This study |
| YqgC-NOS936-LFHF | GTTACTTTTTATGGACGGTCCTATAAATGGCTTACGAACTTCCAGAATTACCTTATG | This study |
| YqgC-NOS936-LFHR | CATAAGGTAATTCTGGAAGTTCGTAAGCCATTTATAGGACCGTCCATAAAAAGTAAC | This study |
| 0.691-yqgC-R | **AAGGCCTTATTGGCC**TAGTCTGTCTCACTCAAAC | This study |
| sodA-FLAG-LFHF | CAAAAGATTATAAAGATGATGATGATAAATAA TGGCACAAACAAGGTCCTCATTATG | This study |
| sodA-FLAG-LFHR | TTATTTATCATCATCATCTTTATAATCTTTTGCTTCGCTGTATAGACGAGCCACTTC | This study |
| 1.571-sodA-R | **AAGGCCTTATTGGCC**ATCGGAAGCGACTTCTTTTTCTCTTTATG | This study |
| yqgC-seqF | GAGGCAGTTTTTGCCGCAGTATTATTC | This study |
| sodA-seqR | TGTCCAGAATAATTTGTGGTTCGCGTG | This study |
| yceE-repair-F | **AAGGCCAACGAGGCC**CAGTATGGCCATTCAATTATCAAAAGG | This study |
| yceG-repair-R | **AAGGCCTTATTGGCC**AACCGAGCTCCTGCTTCGGGTCGAC | This study |
| yceF-206-LFH-Fw | CACAGCATTCGTCTTAATCGGCATTACCGCGCTGAAAATGGCGGGGAGCGCTTTC | This study |
| yceF-206-LFH-Rev | GAAAGCGCTCCCCGCCATTTTCAGCGCGGTAATGCCGATTAAGACGAATGCTGTG | This study |
| yceF-seqF | GCAAAAGTGTTCCTTGTGCTGATTGA | This study |
| yceF-seqR | GTTTTATTCTTCTTTTGAAGCGGCTG | This study |

Sequences shown in red or blue are recognition sites for SfiI used for cloning of repair templates into pAJS23.

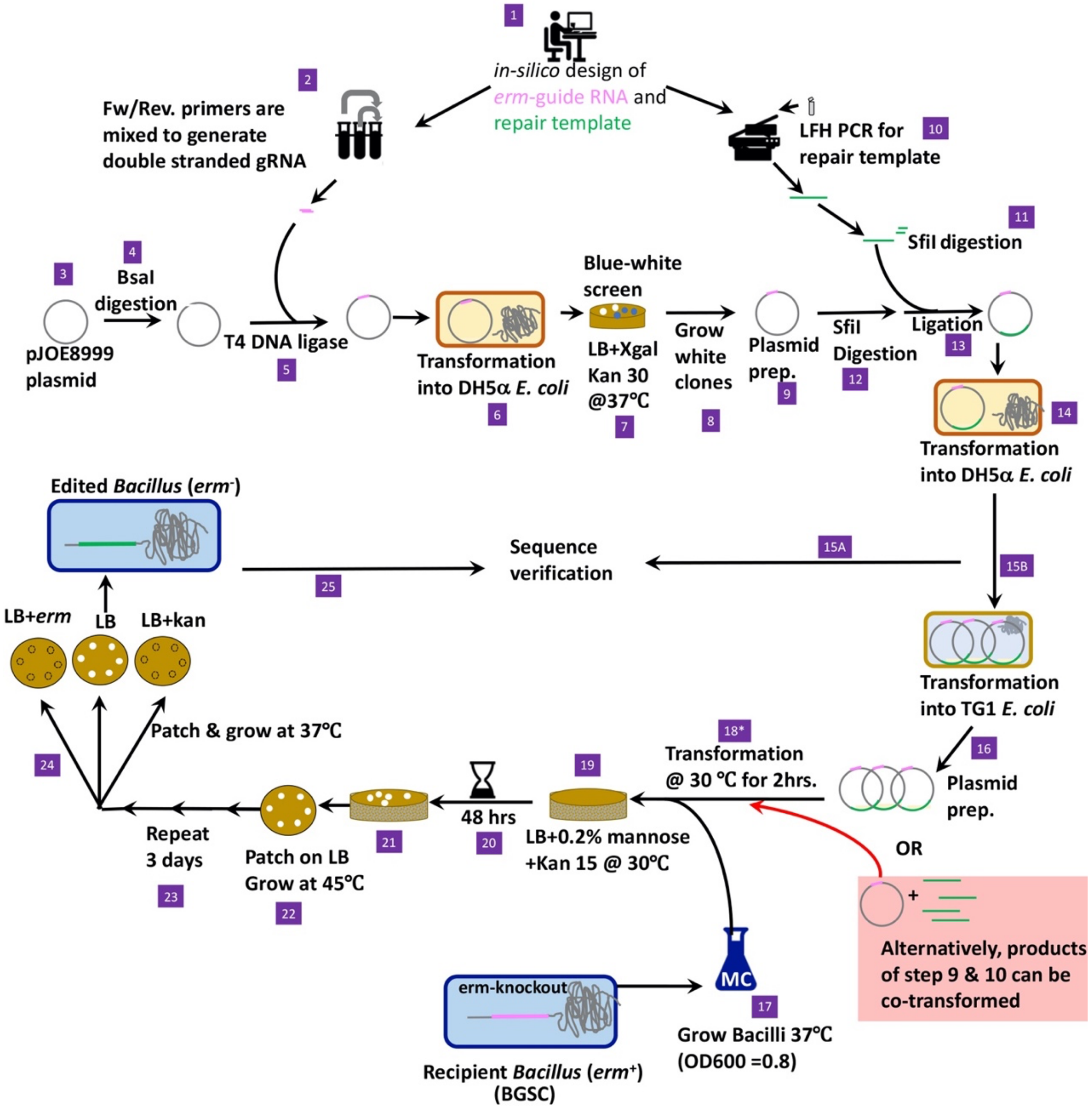


**Fig S1**: **Detailed outline of CRISPR-Cas based editing strategy.** Steps 1-8 shows erm-guide RNA cloning in pJOE8999 (ref. 1) to generate pAJS23 (step 9). For the applications here, begin at step 9 and clone a repair template into pAJS23 at the SfiI sites, or instead provide a suitable repair template in the form of either a PCR product or genomic DNA as shown in pink box at step 18* with red arrow.

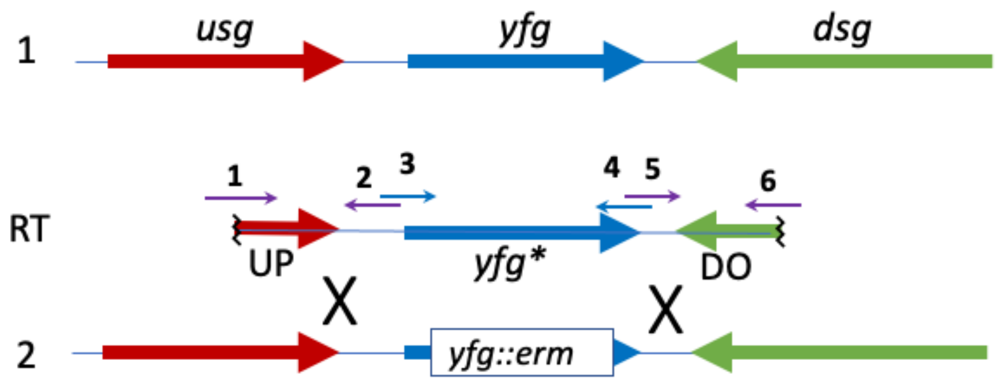


**Fig. S2**. **General strategy for assembly of repair template.** A generic chromosome region (1) is shown, containing your favorite gene (*yfg*) flanked by an upstream gene (*usg*) and a downstream gene (*dsg*). To alter *yfg* (or transfer a different *yfg* allele) into your strain, first introduce the relevant *erm* disruptant (*yfg::erm*) from the BKE collection (2). One can then introduce into strain 2 any genomic DNA containing a desired *yfg** allele by cotransformation with pAJS23 (which will cleave the *erm* cassette). Alternatively, mutations and gene fusions can be constructed using a repair template (RT) constructed by PCR-based methods. The desired cargo fragment (blue primers 3 and 4) can contain altered function alleles, gene fusions, or orthologous genes, for example. This fragment is then linked by SOEing PCR (taking advantage of complementary sequences between divergent primers 2/3 and 4/5, as shown) with an UP (primer 1 and 2) and a DO (primer 5 and 6) fragment for homologous recombination. The UP and DO fragments should be at least ~500 bp for the best efficiency. Note that to clone the RT into pAJS23, primers 1 and 6 should additionally include distinct SfiI recognition sequences. For example, primer 1 can include the 5'-sequence AAGGCCAACGAGGCC and primer 6 can include the 5'-sequence AAGGCCTTATTGGCC (see Table S2 for examples highlighted in red and blue, respectively).

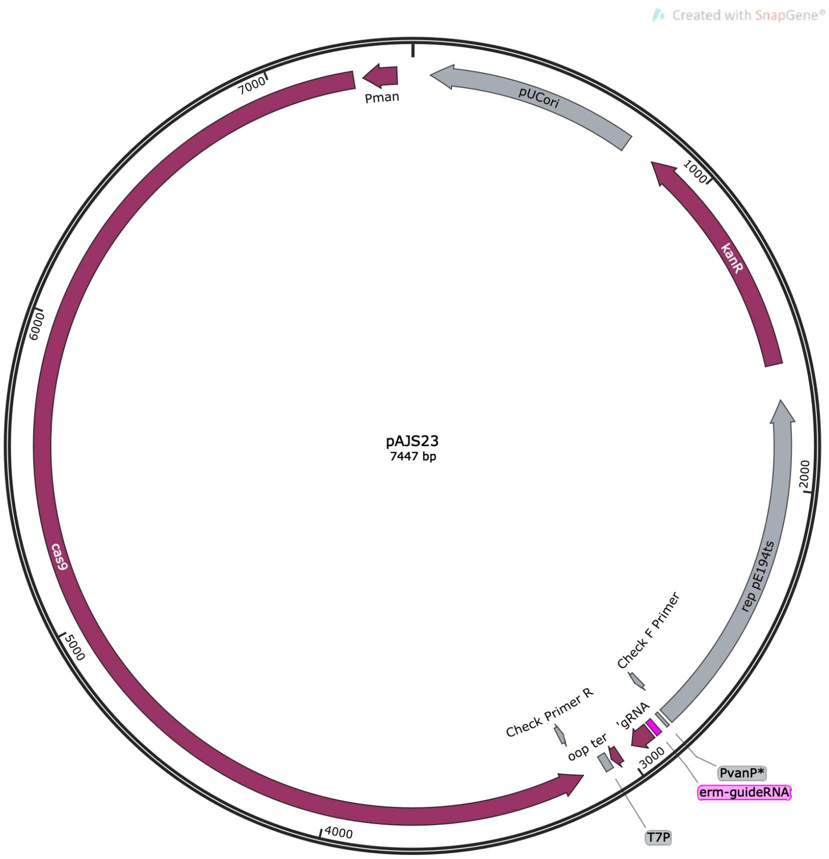


>pAJS23
gaatggccatgaccaaaatcccttaacgtgagttttcgttccactgagcgtcagaccccgtagaaaagatcaaaggatcttcttgagatcctttttttctgcgcgtaatctgctgcttgcaaacaaaaaaaccaccg
ctaccagcggtggtttgtttgccggatcaagagctaccaactctttttccgaaggtaactggcttcagcagagcgcagataccaaatactgtccttctagtgtagccgtagttaggccaccacttcaagaactctgta
gcaccgcctacatacctcgctctgctaatcctgttaccagtggctgctgccagtggcgataagtcgtgtcttaccgggttggactcaagacgatagttaccggataaggcgcagcggtcgggctgaacggggggtt
cgtgcacacagcccagcttggagcgaacgacctacaccgaactgagatacctacagcgtgagctatgagaaagcgccacgcttcccgaagggagaaaggcggacaggtatccggtaagcggcagggtcgga
acaggagagcgcacgagggagcttccagggggaaacgcctggtatctttatagtcctgtcgggtttcgccacctctgacttgagcgtcgatttttgtgatgctcgtcaggggggcggagcctatggaaaaacgcc
agcaacgcggcctttttacggttcctggccttttgctggccttttgctcacatgttctttcctgcgttatcccctgattctgtggataaccgtattaccgcctttgagtgagctgaaattatgaggggatctctcagagct
cgaggtcatcgttcaaaatggtatgcgttttgacacatccactatatatccgtgtcgttctgtccactcctgaatcccattccagaaattctctagcgattccagaagtttctcagagtcggaaagttgaccagacatt
acgaactggcacagatggtcataacctgaaggaagatctgattgcttaactgcttcagttaagaccgaagcgctcgtcgtataacagatgcgatgatgcagaccaatcaacatggcacctgccattgctacctgc
acagtcaaggatggtagaaatgttgtcggtccttgcacacgaatattacgccatttgcctgcatattcaaacagctcttctacgataagggcacaaatcgcatcgtggaacgtttgggcttctaccgatttagcagtt
tgatacactttctctaagtatccacctgaatcataaatcggcaaaatagagaaaaattgaccatgtgtaagcggccaatctgattccacctgagatgcataatctagtagaatctcttcgctatcaaaattcacttc
caccttccactcaccggttgtccattcatggctgaactctgcttcctctgttgacatgacacacatcatctcaatatccgaatagggcccatcagtctgacgaccaagagagccataaacaccaatagccttaacat
catccccatatttatccaatattcgttccttaatttcatgaacaatcttcattctttcttctctagtcattattattggtccattcactattctcattcccttttcagataattttagatttgcttttctaaataagaatatttgg
agagcaccgttcttattcagctattaaacccattatatcgggtttttgaggggatttcaactgcagacacctaaattcaaaatctatcggtcagatttataccgatttgattttatatattcttgaataacatacgccga
gttatcacataaaagcgggaaccaatcatcaaatttaaacttcattgcataatccattaaactcttaaattctacgattccttgttcatcaataaactcaatcatttctttaattaatttatatctatctgttgttgttttc
tttaataattcatcaacatctacaccgccataaactatcatatcttctttttgatatttaaatttattaggatcgtccatgtgaagcatatatctcacaagacctttcacacttcctgcaatctgcggaatagtcgcattc
aattcttctgttaattatttttatctgttcataagatttattaccctcatacatcactagaatatgataatgctctttttttcatcctaccttctgtatcagtatccctatcatgtaatggagacactacaaattgaatgtgta
actcttttaaatactctaaccactcggcttttgctgattctggatataaaacaaatgtccaattacgtcctcttgaatttttcttgttttcagtttcttttattacattttcgctcatgatataataacggtgctaatacattt
aacaaaatttagtcatagataggcagcatgccagtgctgtctatcttttttttgtttaaaatgcaccgtattcctcctttgcatatttttttattagaataccggttgcatctgatttgctaatattatatttttctttgattct
atttaatatctcattttcttctgttgtaagtcttaaagtaacagcaactttttttctcttcttttctatctacaaccatcactgtacctcccaacatctgtttttttcactttaacataaaaaacaacctttttaacattaaaaa
cccaatatttatttatttgtttggacaatggacaatggacacctagggggggaggtcgtagtacccccctatgttttctcccctaaataaccccaaaaatctaagaaaaaaagacctcaaaaaggtctttaattaaca
tctcaaatttcgcatttattccaatttcctttttgcgtgtgatgcgaattcttgaccgtgattagagaattgagtaaaatgtacctacgTTTGAAATCGGCTCAGGAAAgttttagagctagaaatagcaagt
taaaataaggctagtccgttatcaacttgaaaaagtggcaccgagtcggtgctttttactccatctggatttgttcagaacgctcggttgccgccgggcgtttttttatctaaagcttaggcccagtcgaaagactggg
cctttttaatacgactcactatagggtcgacggccaacgaggcccgggccaataaggcctttctagattaagaaataatcttcatctaaaatatacttcagtcacctcctagctgactcaaatcaatgcgtgtttcat
aaagaccagtgatggattgatggataagagtggcatctaaaacttcttttgtagacgtatatcgtttacgatcaattgttgtatcaaaatatttaaaagcagcgggagctccaagattcgtcaacgtaaataaatg
aataatattttctgcttgttcacgtattggtttgtctctatgtttgttatatgcactaagaactttatctaaattggcatctgctaaaataacacgcttagaaaattcactgatttgctcaataatctcatctaaataatgc
ttatgctgctccacaaacaattgtttttgttcgttatcttctggactacccttcaacttttcataatgactagctaaatataaaaaattcacatatttgcttggcagagccagctcatttcctttttgtaattctccggcac
tagccagcatccgtttacgaccgttttctaactcaaaaagactatatttaggtagtttaatgattaagtctttttttaacttccttatatcctttagcttctaaaaagtcaatcggatttttttcaaaggaacttctttccat
aattgtgatccctagtaactctttaacggattttaacttcttcgatttcccttttttccaccttagcaaccactaggactgaataagctaccgttggactatcaaaaccaccatattttttggatcccagtctttttttacga
gcaataagcttgtccgaatttctttttggtaaaattgactccttggagaatccgcctgtctgtacttctgttttcttgacaatattgacttggggcatggacaatactttgcgcactgtggcaaaatctcgccctttatcc
cagacaatttctccagtttccccattagtttcgattagagggcgtttgcgaatctctccatttgcaagtgtaatttctgttttgaagaagttcatgatattagagtaaaagaaatattttgcggttgctttgcctatttctt
gctcagacttagcaatcattttacgaacatcataaactttataatcaccatagacaaactccgattcaagttttggatatttcttaatcaaagcagttccaacgacggcatttagatacgcatcatgggcatgatggt

aattgttaatctcacgtactttatagaattggaaatcttttcggaagtcagaaactaatttagattttaaggtaatcactttaacctctcgaataagtttatcattttcatcgtatttagtattcatgcgactatccaaaa
tttgtgccacatgcttagtgatttggcgagtttcaaccaattggcgtttgataaaaccagctttatcaagttcactcaaacctccacgttcagctttcgttaaattatcaaacttacgttgagtgattaacttggcgttt
agaagttgtctccaatagtttttcatctttttgactacttcttcacttggaacgttatccgatttaccacgatttttatcagaacgcgttaagaccttattgtctattgaatcgtctttaaggaaactttgtggaacaatgt
gatcgacatcataatcacttaaacgattaatatctaattcttggtccacatacatgtctcttccattttggagataatagagatagagcttttcattttgcaattgagtattttcaacaggatgctctttaagaatctga
cttcctaattctttgataccttcttcgattcgtttcatacgctctcgcgaatttttctggcccttttgagttgtctgattttcacgtgccatttcaataacgatattttctggcttatgccgccccattactttgaccaattcat
caacaacttttacagtctgtaaaatacctttttaatagcagggctaccagctaaatttgcaatatgttcatgtaaactatcgccttgtccagacacttgtgctttttgaatgtcttctttaaatgtcaaactatcatcat
ggatcagctgcataaaattgcgattggcaaaaccatctgatttcaaaaaatctaatattgttttgccagattgcttatccctaataccattaatcaattttcgagacaaacgtccccaaccagtataacggcgacgtt
taagctgtttcatcaccttatcatcaaagaggtgagcatatgttttaagtctttcctcaatcatctccctatcttcaaataaggtcaatgttaaaacaatatcctctaagatatcttcattttcttcattatccaaaaaat
ctttatctttaataatttttagcaaatcatggtaggtacctaatgaagcattaaatctatcttcaactcctgaaatttcaacactatcaaaacattctatttttttgaaataatcttcttttaattgcttaacggttactttt
cgatttgttttgaagagtaaatcaacaatggctttcttctgttcacctgaaagaaatgctggttttcgcattccttcagtaacatatttgacctttgtcaattcgttataaaccgtaaaatactcataaagcaaactatg
ttttggtagtactttttcatttggaagatttttatcaaagtttgtcatgcgttcaataaatgattgagctgaagcacctttatcgacaacttcttcaaaattccatggggtaattgtttcttcagacttccgagtcatccat
gcaaaacgactattgccacgcgccaatggaccaacataataaggaattcgaaaagtcaagatttttttcaatcttctcacgattgtcttttaaaaatggataaaagtcttcttgtcttctcaaaatagcatgcagctc
acccaagtgaatttgatggggaatagagccgttgtcaaaggtccgttgcttgcgcagcaaatcttcacgatttagtttcaccaataattcctcagtaccatccatttttttctaaaattggtttgataaatttataaaatt
cttcttggctagctcccccatcaatataacctgcatatccgtttttttgattgatcaaaaaagatttctttatacttttctggaagttgttgtcgaactaaagcttttaaaagagtcaagtcttgatgatgttcatcgtagc
gtttaatcattgaagctgataggggagccttagttatttcagtatttactcttaggatatctgaaagtaaaatagcatctgataaattcttagctgccaaaaacaaatcagcatattgatctccaatttgcgccaata
aattatctaaatcatcatcgtaagtatcttttgaaagctgtaatttagcatcttctgccaaatcaaaatttgatttaaaattaggggtcaaacccaatgacaaagcaatgagattcccaaataagccatttttcttctc
accggggagctgagcaatgagattttctaatcgtcttgatttactcaatcgtgcagaaagaatcgctttagcatctactccacttgcgttaatagggttttcttcaaataattgattgtaggtttgtaccaactggata
aatagtttgtccacatcactattatcaggatttaaatctccctcaatcaaaaaatgaccacgaaacttaatcatatgcgctaaggccaaatagattaagcgcaaatccgctttatcagtagaatctaccaattttttt
cgcagatgatagatagttggatatttctcatgataagcaacttcatctactatatttccaaaaataggatgacgttcatgcttcttgtcttcttccaccaaaaaaagactcttcaagtcgatgaaagaaactatcatct
actttcgccatctcatttgaaaaaatctcctgtagataacaaatacgattcttccgacgtgtataccttctacgagctgtccgtttgagacgagtcgcttccgctgtctctccactgtcaaataaaagagcccctataa
gatttttttgatactgtggcggtctgtatttcccagaaccttgaactttttagacggaaccttataatcatcagtgatcaccgcccatccgacgctatttgtgccgatatctaagcctattgagtatttcttatccatttt
tgcctcctaacttaagaataagatcttgtctcaactgtataccgaaatcagctcattaaaatcgctttttttaccataggttccggtaataaaggcatttttccctataacaaaaaaagcaaggaataatccctgcttt
taataatccaaatgagataaaaatgtcatgacattggtgtacagaa

**Fig. S3**. **Plasmid map and DNA sequence of plasmid pAJS23.** A plasmid map of the 7447 bp pAJS23 plasmid (derived from pJOE8999, ref. 1) is shown illustrating key features. The text file contains the complete plasmid sequence, with the sequence corresponding to the anti-*erm* gRNA in uppercase.

### Bibliography and References Cited


1. Altenbuchner J. 2016. Editing of the Bacillus subtilis Genome by the CRISPR-Cas9 System. Appl Environ Microbiol 82:5421-7.
